# Remdesivir potently inhibits SARS-CoV-2 in human lung cells and chimeric SARS-CoV expressing the SARS-CoV-2 RNA polymerase in mice

**DOI:** 10.1101/2020.04.27.064279

**Authors:** Andrea J. Pruijssers, Amelia S. George, Alexandra Schäfer, Sarah R. Leist, Lisa E. Gralinksi, Kenneth H. Dinnon, Boyd L. Yount, Maria L. Agostini, Laura J. Stevens, James D. Chappell, Xiaotao Lu, Tia M. Hughes, Kendra Gully, David R. Martinez, Ariane J. Brown, Rachel L. Graham, Jason K. Perry, Venice Du Pont, Jared Pitts, Bin Ma, Darius Babusis, Eisuke Murakami, Joy Y. Feng, John P. Bilello, Danielle P. Porter, Tomas Cihlar, Ralph S. Baric, Mark R. Denison, Timothy P. Sheahan

## Abstract

Severe acute respiratory syndrome coronavirus 2 (SARS-CoV-2) emerged in 2019 as the causative agent of the novel pandemic viral disease COVID-19. With no approved therapies, this pandemic illustrates the urgent need for safe, broad-spectrum antiviral countermeasures against SARS-CoV-2 and future emerging CoVs. We report that remdesivir (RDV), a monophosphoramidate prodrug of an adenosine analog, potently inhibits SARS-CoV-2 replication in human lung cells and primary human airway epithelial cultures (EC_50_ = 0.01 μM). Weaker activity was observed in Vero E6 cells (EC_50_ = 1.65 μM) due to their low capacity to metabolize RDV. To rapidly evaluate *in vivo* efficacy, we engineered a chimeric SARS-CoV encoding the viral target of RDV, the RNA-dependent RNA polymerase, of SARS-CoV-2. In mice infected with chimeric virus, therapeutic RDV administration diminished lung viral load and improved pulmonary function as compared to vehicle treated animals. These data provide evidence that RDV is potently active against SARS-CoV-2 *in vitro* and *in vivo*, supporting its further clinical testing for treatment of COVID-19.

## INTRODUCTION

Coronaviruses (CoV) are genetically diverse positive sense RNA viruses that circulate in animals and humans. In the past 20 years, three new human CoV have emerged: severe acute respiratory syndrome CoV (SARS-CoV-1) in 2002, Middle East respiratory syndrome (MERS)-CoV in 2012 and current pandemic SARS-CoV-2, the causative agent of COVID-19 (de Wit et al., 2016; Zhou et al., 2020b). While four endemic human CoV (HCoV-OC43, −229E, -NL63, and -HKU1) typically cause mild respiratory diseases with common cold-like symptoms, SARS-CoV-1, MERS-CoV, and SARS-CoV-2 cause severe respiratory disease with respective mortality rates of 11% (Chan-Yeung and Xu, 2003), 35% (Arabi et al., 2017), and an estimated 3% (Chen, 2020). The development of effective broad-spectrum antivirals has been hampered by viral diversity, the capacity of CoVs to adaptively overcome negative selective pressures, and the ability to actively counteract drugs through the action of a proofreading exoribonuclease. We previously reported that remdesivir (RDV), a monophosphoramidate prodrug of the *C*-adenosine analog GS-441524, potently inhibits replication of a broad spectrum of pre-pandemic bat CoVs and human epidemic CoVs in primary human lung cell cultures (Agostini et al., 2018; Brown et al., 2019; Sheahan et al., 2017). Biochemical analysis of the mechanism of inhibition of the SARS-CoV-2, SARS-CoV-1, and MERS-CoV RNA-dependent RNA polymerase (RdRp) revealed that incorporation of the active metabolite RDV triphosphate (RDV-TP) was more efficient than the natural substrate ATP and led to delayed chain termination three nucleotides downstream of incorporation (Gordon et al., 2020a, 2020b). Prolonged passaging of murine hepatitis virus (MHV), a group 2a CoV, in the presence of GS-441524 resulted in low level resistance through mutations in the RdRp, further implicating this protein as the drug target (Agostini et al., 2018). RDV showed both prophylactic and therapeutic efficacy in mouse models of SARS and MERS and against MERS-CoV challenge in a rhesus macaque model (Sheahan et al., 2017, 2020a; Wit et al., 2020). Here we report that RDV potently inhibits SARS-CoV-2 replication Calu3 human lung cells with sub-micromolar EC_50_ and in primary human airway epithelial cultures (HAEs) with nanomolar EC_50_. Notably, we have detected comparably lower potency of RDV in established human and monkey cell lines due to their lower metabolic capacity to activate the compound. Mice infected with chimeric SARS-CoV-1 encoding the SARS-CoV-2 RdRp and treated therapeutically with RDV show decreased viral loads in the lungs and increased pulmonary function. These data emphasize the potential of RDV as a promising countermeasure against the ongoing COVID-19 pandemic.

## RESULTS

### Structural model of remdesivir incorporation by the SARS-CoV-2 polymerase and conservation of the active site across human CoV

Drug function and performance is heavily influenced by microvariation in target genes across virus families, biodistribution in the organism, and, importantly, host cell and tissue expression patterns that influence drug stability and metabolism. We previously modeled RDV on a homology model of SARS-CoV-2 based on the cryo-EM structure of SARS-CoV-1 polymerase complex (Gordon et al., 2020b; Kirchdoerfer and Ward, 2019). Composed of nsp12, nsp7 and nsp8, the model was consistent with biochemical findings predicting efficient incorporation of RDV-TP into the growing RNA strand and provided an explanation for the observed delayed chain termination after the incorporation of three additional nucleotides. We have since refined this model using the recently released cryo-EM structure of the SARS-CoV-2 polymerase complex (Gao et al., 2020). The major qualitative change is a more complete picture of the N-terminal NiRAN domain of nsp12, which was not resolved in the SARS-CoV-1 structure. The current model of the pre-incorporation state, with RDV-TP, RNA primer and template strands and catalytic metals was well-optimized with a series of constrained energy minimizations and conformational searches, as described previously (**Fig. 1A**). Bound to the two catalytic Mg^2+^ ions, RDV-TP is coordinated by two basic residues (R553 and R555). The ribose 2’OH forms hydrogen bonds to T680 and N691, and the 1’CN resides in a shallow pocket formed by T687 and A688 (**Fig. 1B**). The interaction with T680 distinguishes CoVs from other structurally related families, including noroviruses, picornaviruses, and the flaviviruses. While key residues including D623, S682, and N691, are conserved across all these virus families and have been shown to govern positioning of the NTP into the active site, the role of T680 appears to be novel. While writing this manuscript, another model of RDV-TP in the SARS-CoV-2 active site (Shannon et al., 2020) was published which predicts a role for S682 as well. The position of T680 relative to N691 strongly implies it will contribute to the recognition of the ribose 2’OH, likely diminishing the role of S682 as a result, consistent with earlier predictions (Kirchdoerfer and Ward, 2019).

**Figure 1.**
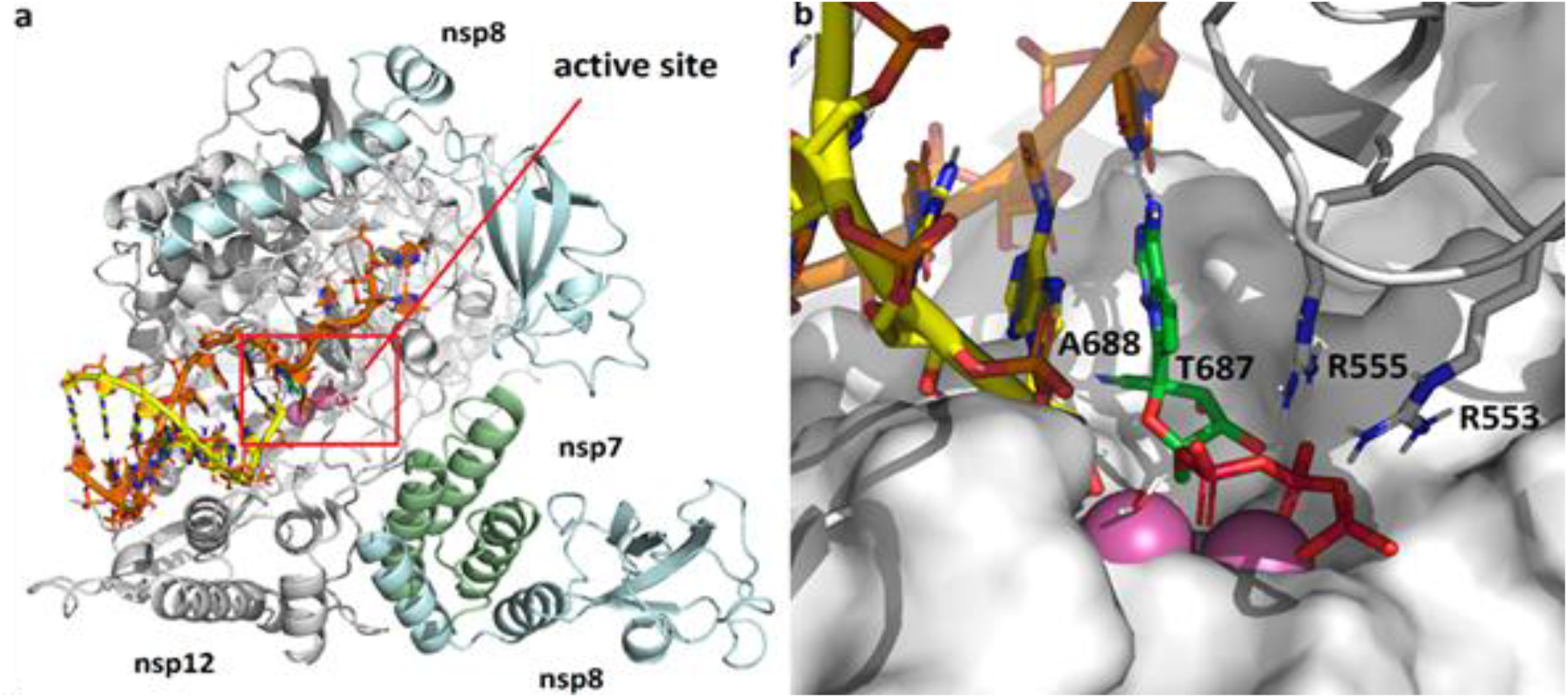
Modeling of remdesivir onto the SARS-CoV-2 RdRp structure. **A.** Model of SARS-CoV-2 polymerization complex in its elongating state. The model was based on the cryo-EM apo structures of SARS-CoV-1 (PDB 6NUR) and SARS-CoV-2 (PDB 6M71). The active site is boxed in red. **B.** Enlarged view of active site depicting RDV pre-incorporation. The 1’CN substituent sits in a shallow pocket formed by residues T687 and A688. Bound to the two catalytic Mg^2+^ ions (pink), the triphosphate is coordinated by two basic residues (R553 and R555)

Modeling of the RDV resistance mutations identified in MHV (Agostini et al., 2018) onto homologous residues V557 and F480 in the SARS-CoV-2 RdRp structure reveals that V557L shifts the position of the template base, which in turn shifts the positioning of the incoming NTP (**Fig. S1 A, B**). This will impact RDV activity in that it alters the position of the 1’CN in the pocket. Because the model predicts no direct interaction of F480 with the NTP, primer, or template, the effect of the F480L mutation is more difficult to discern. The F480L mutation could potentially induce a subtle change to the 1’CN binding pocket (**Fig. S1 C, D**). Alignment of nsp12 sequences from SARS-CoV-2 used in other studies of RDV shows complete conservation of nsp12 nucleotide sequences, predicting the comparable antiviral activity of RDV against these isolates (**Fig. S2**). We next modeled the active sites of the six other human CoVs SARS-CoV-1 (**Fig. S3A**), MERS-CoV (**Fig. S3B**), HCoV-OC43 (**Fig. S3C)**, −229E (**Fig. S3D**), -NL63 (**Fig. S3E**), and -HKU1 (**Fig. S3F**). The models show that SARS-CoV-2 is identical to SARS-CoV-1 out to a radius of 18 Å from the active site. Differences detected on the periphery of the active site of the MERS-CoV and HCoV-OC43, −229E, -NL63, and HKU1 correspond to residues that do not directly interact RDV-TP. Together, these data demonstrate high structural conservation of the RdRp active site interacting with RDV-TP across all seven known human CoV strains.

### Remdesivir and GS-441524 potently inhibit SARS-CoV-2 replication

Remdesivir (RDV) and its parent nucleoside analog GS-441524 inhibit CoVs and multiple other viruses (Agostini et al., 2018; Cho et al., 2012; Lo et al., 2017; Sheahan et al., 2017; Warren et al., 2016). Previous reports (Choy et al., 2020; Wang et al., 2020; Runfeng et al., 2020) suggest RDV inhibits SARS-CoV-2, but comparative studies of anti-SARS-CoV-2 activity using authentic compound in physiologically relevant cell lines are lacking. We first compared SARS-CoV-2 replication in established cell lines to determine which cell types could potentially be suitable for studying RDV efficacy against SARS-CoV-2. Viral yields were determined at 24, 48, and 72 hours post-infection (hpi) in Vero E6, Vero CCL-81 (Vero), Huh7, and Calu3 2B4 (Yoshikawa et al., 2010) cells (**Fig. 2A**). Vero E6 and Vero cells supported highest levels of SARS-CoV-2 replication, consistent with a previous study (Harcourt et al.). Maximum yields were detected at 48 hpi in Vero E6 cells (>6 logs at MOI = 0.1 and 0.01 PFU/cell), 24 hpi in Vero cells infected at MOI = 0.1 PFU (>5 logs), 48 hpi in Vero cells infected at MOI = 0.01 PFU/cell (>5 logs), 72 hpi in Calu3 2B4 (>4 logs at MOI = 0.1 PFU/cell), and 48 hpi in Huh7 cells (>4 logs at MOI = 0.1 PFU/cell, <2 logs at MOI = 0.01 PFU/cell). These results indicate that Vero E6, Vero, and Calu3 2B4 cells support varying levels of SARS-CoV-2 replication and cell type should be chosen for a given study depending on study goals.

**Figure 2.**
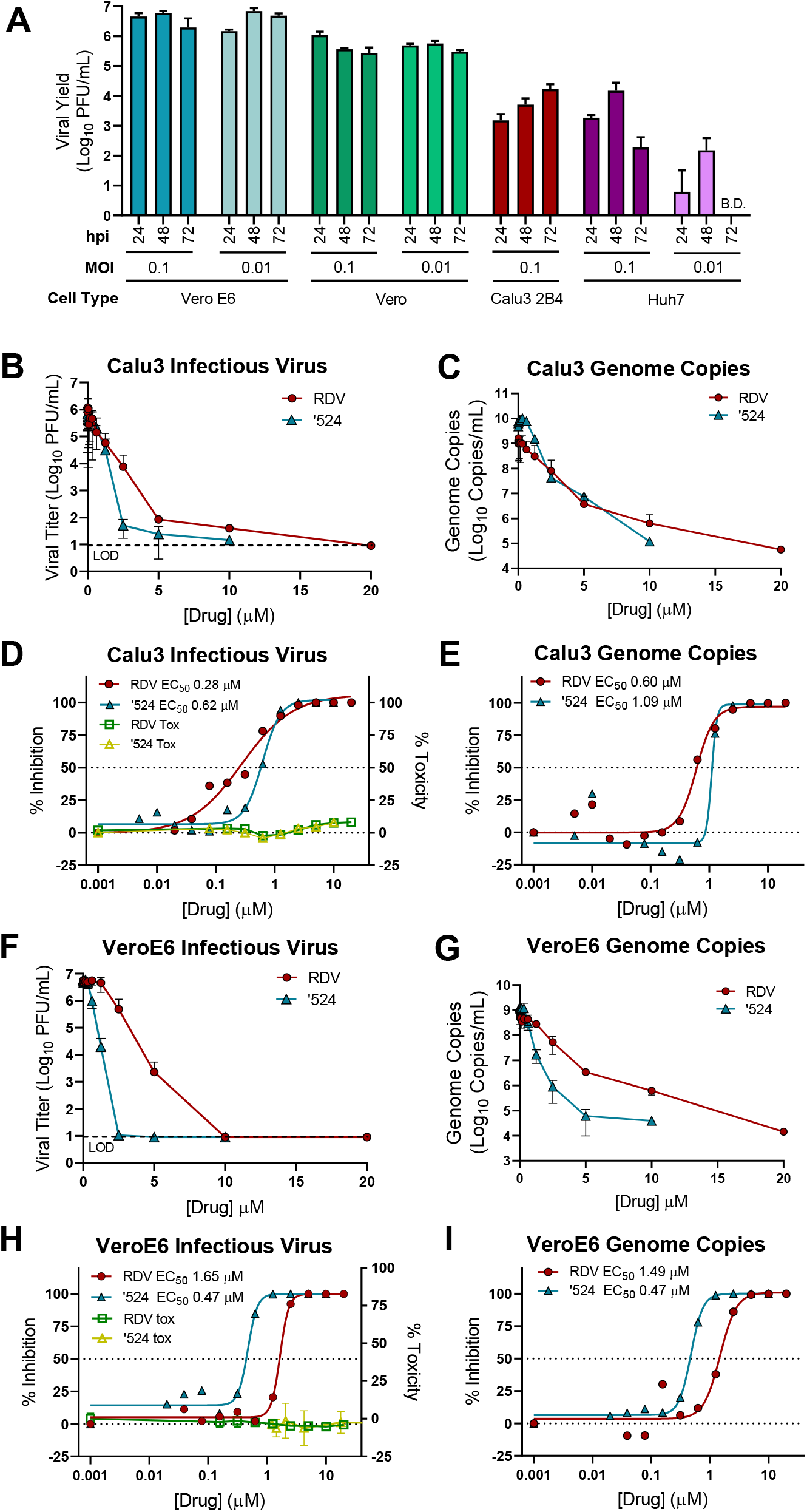
Prodrug remdesivir (RDV) and parent nucleoside GS-441524 (‘524) potently inhibit SARS-CoV-2 replication. **A.** Vero E6, Vero CCL-81 (Vero), Huh7, and Calu3 2B4 cells were infected with MOI = 0.01 and/or 0.1 PFU/cell SARS-CoV-2 (2019-nCoV/USA-WA1/2020), and infectious viral titers were determined by plaque assay at 0.5, 24, 48, and 72 hours post-infection (hpi). Viral yields were calculated by subtracting the average 0.5 h (post-adsorption, pre-incubation) titer from each subsequent time point. Data represent the average of three replicates from one experiment. Error bars indicate SD. B.D.: below detection. Calu3 cells were infected with 0.1 PFU/cell SARS-CoV-2 and Vero cells were infected with 0.01 PFU/cell SARS-CoV-2 and treated with RDV, GS-441524 (‘524), or DMSO only (control) in cell culture medium. Supernatants were collected at 48 h (Vero E6) or 72 h (Calu3) post-infection. **B, C.** Reduction of SARS-CoV-2 replication by RDV in Calu3 cells as determined by infectious viral titer and RT-qPCR. **D**. Percent inhibition of SARS-CoV-2 replication by RDV and GS-441524 in Calu3 as determined by infectious viral titer [RDV: EC_50_ = 0.28 μM, EC_90_ = 2.48 μM; GS-441524 EC_50_ = 0.62 μM, EC_90_ = 1.34 μM]. No significant cytotoxicity of either compound was detected in Calu3 cells. **E.** Percent inhibition of SARS-CoV-2 replication by RDV and GS-441524 in Calu3 as determined RT-qPCR [RDV: EC_50_ = 0.60 μM, EC_90_ = 1.28 μM; GS-441524: EC_50_ = 1.09 μM, EC_90_ = 1.37 μM]. **F, G.** Reduction of SARS-CoV-2 replication by RDV in Vero E6 cells as determined by infectious viral titer and RT-qPCR. **H.** Percent inhibition of SARS-CoV-2 replication by RDV and GS-441524 in Vero E6 cells as determined by infectious viral titer [RDV: EC_50_ = 1.65 μM, EC_90_ = 2.40 μM; GS-441524: EC_50_ = 0.47 μM, EC_90_ = 0.71 μM]. No significant cytotoxicity of either compound was detected in Vero E6 cells. **I.** Percent inhibition of SARS-CoV-2 replication by RDV and GS-441524 in Vero E6 cells as determined RT-qPCR [RDV: EC_50_ = 1.49 μM, EC_90_ = 3.03 μM; GS-441524: EC_50_ = 0.47 μM, EC_90_ = 0.80 μM]. Data represent means of 2-4 independent experiments with 2-3 replicated each. Error bars represent SEM.

To determine if RDV and GS-441524 inhibit SARS-CoV-2 replication in established cell lines, Calu3 2B4 human lung adenocarcinoma cells and Vero E6 African green monkey kidney cells were infected with the SARS-CoV-2 clinical isolate 2019-nCoV/USA-WA1/2020 and treated with a range of RDV or GS-441524 concentrations. Supernatants were harvested at time points corresponding to peak viral replication for each cell type, and infectious viral titer and viral genome copy number in the supernatant were quantified by plaque assay and RT-qPCR, respectively. RDV and GS-441524 potently inhibited SARS-CoV-2 replication in a dose-dependent manner in both cell types (**Fig. 2; Table 1**). In Calu3 cells, both compounds displayed dose-dependent inhibition of viral replication as determined by plaque assay (**Fig. 2B**) and RT-qPCR (**Fig. 2C**). RDV inhibited SARS-CoV-2 with an EC_50_ = 0.28 μM and EC_90_ = 2.48 μM. The parent compound GS-441524 was less potent: EC_50_ = 0.62 μM, EC_90_ = 1.34 μM (**Fig. 2D; Table 1**). EC_50_ values determined by quantification of viral genome copies were roughly two-fold higher than those obtained by quantification of infectious virus (**Fig. 2E; Table 1**). Both compounds also displayed dose-dependent inhibition of viral replication in Vero E6 cells as determined by infectious viral titer and genome copy number (**Fig 2F)**. RDV inhibited SARS-CoV-2 with EC_50_ = 1.65 μM and EC_90_ = 2.40 μM, while GS-441524 was more potent (EC_50_ = 0.47 μM, EC_90_ = 0.71 μM) (**Fig. 2G; Table 1**). Relative potency based on genome copies was similar to that assessed by quantification of infectious viral titer in Vero E6 cells (**Fig. 2H; Table 1**). Thus, RDV inhibits SARS-CoV-2 more potently in Calu3 2B4 than in Vero E6 cells.

**Table 1:** Cell-specific SARS-CoV-2 potency and metabolism.

| Analysis | RDV |  |  | GS-441524<sup>a</sup> |  |
| --- | --- | --- | --- | --- | --- |
|  | Vero E6 | Calu3 2B4 | HAE<sup>b</sup> | Vero E6 | Calu3 2B4 |
| Plaque assay<br>EC<sub>50</sub> (μM) | 1.65 | 0.28 | 0.010 | 0.47 | 0.62 |
| Genome copy<br>EC<sub>50</sub> (μM) | 1.49 | 0.60 | n.d. | 0.47 | 1.09 |
| RDV-TP at 24 h<sup>c</sup><br>(pmol/million cells) | 0.54 ± 0.15<sup>d</sup> | 2.17 ± 0.14<sup>d</sup> | 10.6 ± 5.3<sup>b</sup> | 2.37 ± 0.22<sup>d</sup> | 0.85 ± 0.16<sup>d</sup> |

<sup>a</sup> GS-441524 was not tested for antiviral potency nor RDV-TP levels in HAE cultures
<sup>b</sup> HAE antiviral potencies and RDV-TP levels were determined independently in differentiated cultures from two donors. RDV-TP levels in HAEs are presented as the mean ± SD of quadruplicate technical replicates from two donors
<sup>c</sup> Individual analyte data are presented in the supplementary information
<sup>d</sup> Values represent mean ± SD from two independent experiments with each performed with duplicate samples

### RDV is a highly potent antiviral inhibitor of SARS-CoV-2 in primary human airway epithelial (HAE) cultures

Primary HAE cultures grown on air-liquid interface recapitulate the cellular complexity and physiology of the human conducting airway (Sims et al., 2005). Therefore, we evaluated antiviral activity of RDV in this biologically relevant model. In RDV treated HAE, we observed a dose-dependent reduction in infectious virus production, with >100-fold inhibition at the highest tested concentration (**Fig. 3A**). Importantly, RDV demonstrate potent antiviral activity with EC_50_ values of 0.0010 and 0.009 μM in two independent experiments (**Fig. 3B**). We previously reported that RDV is not cytotoxic at doses at or below 10 μM in this culture system, supporting the conclusion that the observed antiviral effect was virus-specific (Sheahan et al., 2017). Together, these data demonstrate that RDV is potently antiviral against SARS-CoV-2 in primary human lung cultures with a selectivity index of >1000.

**Figure 3.**
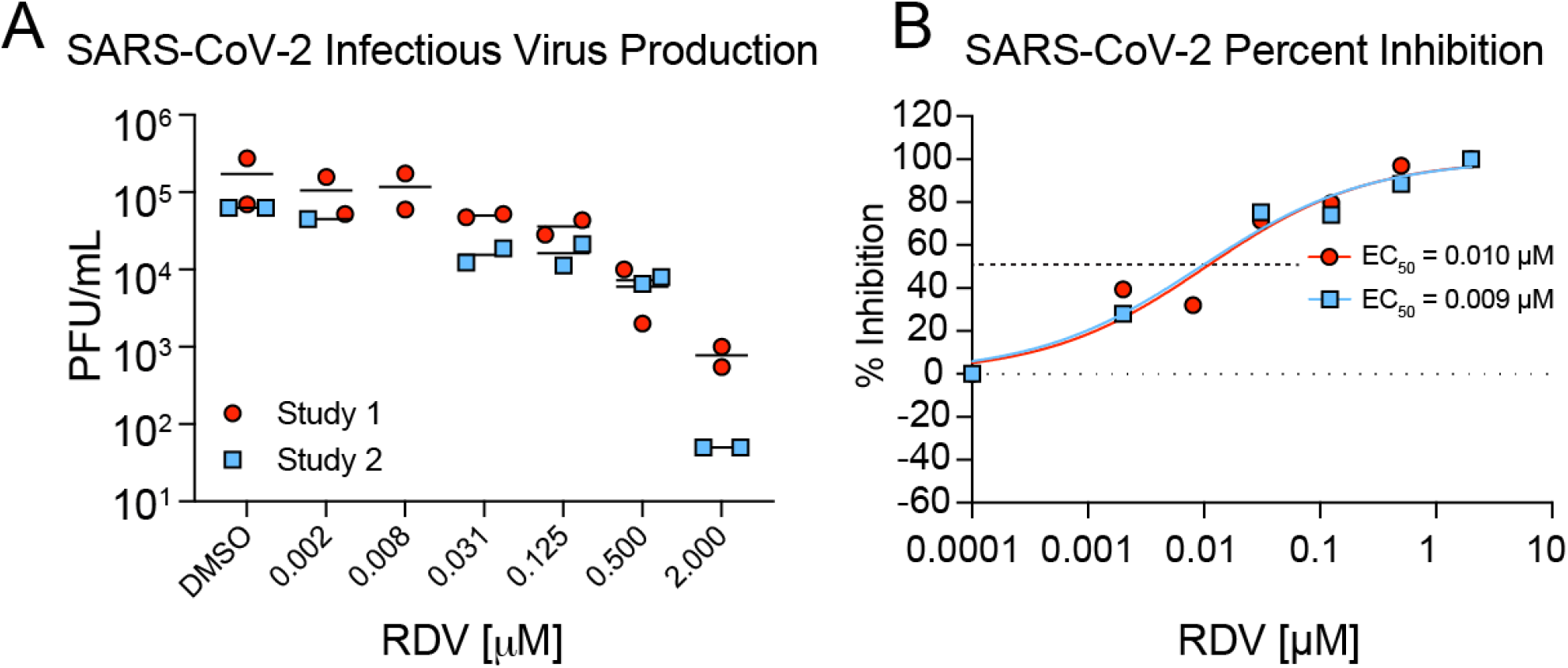
RDV is potently antiviral against SARS-CoV-2 in primary human airway epithelial (HAE) cultures. HAE cultures were infected with a SARS-CoV-2 clinical isolate (2019-nCoV/USA-WA1/2020) at MOI = 0.5 PFU/cell for 2 h, after which virus was removed and cultures were washed 3 times, followed by incubation 37° C for 48 h. **A.** SARS-CoV-2 infectious virus production in two independent studies. Virus was titered via plaque assay in apical washes at 48 h post-infection. Each symbol represents the titer from a single culture, and line is drawn at the mean. **B.** Percent inhibition generated from titer data in A.

### Antiviral activities of RDV and GS-441524 correlate with RDV-TP metabolite levels

Cell type specific expression of genes that metabolize ribonucleoside analogs can have a profound impact on activity (Eriksson, 2013; Koczor et al., 2012). Table 1 and prior studies (Bojkova et al., 2020; Choy et al., 2020; Jeon et al., 2020) demonstrate the antiviral activity of RDV against SARS-CoV-2 is highly variable in different cell culture models. Both RDV and GS-441524 undergo intracellular conversion to the active metabolite RDV-TP involving several metabolic steps (**Fig. S4**) and the efficiency of each step might differ between cell types. Therefore, to reconcile the differences in antiviral activity of RDV and GS-441524 observed in our and other studies, we compared intracellular RDV-TP concentrations in Vero E6, Calu3 2B4, and HAEs following incubation with the two compounds. RDV-TP levels per million cells produced after 8- to 48-hour treatment with RDV were substantially higher in primary HAE cultures than either Calu3 2B4 or Vero E6. (**Fig 4; Table 1; Tables S1, S2**). Given the primary nature of HAE cultures, we used cells from two independent donors with similar demographic profiles. RDV-TP was efficiently formed in both donor cultures following incubation with RDV with a difference of < 3-fold from each other. The lowest levels of RDV-TP were observed following RDV treatment of Vero E6 cells and were approximately 4- and 20-fold lower than those observed in Calu3 2B4 and HAE cultures, respectively. The levels of GS-441524 as well as the intermediate mono- and di-phosphorylated metabolites (RDV-MP and RDV-DP) were readily detected in Calu3 2B4 cultures following treatment with RDV, but were below the limit of quantification in Vero E6 cells at all time points tested (**Table S1**). In addition, incubation of Vero E6 cells with GS-441524 yielded 4-fold higher RDV-TP levels compared to incubation with RDV corresponding to higher antiviral potency of GS-441524 relative to RDV, which is not observed with either Calu3 or HAE cultures. (**Table S1, S2**). In conclusion, the RDV-TP levels in the different cell types directly correlated with the antiviral potencies of RDV and GS-441524 against SARS-CoV-2 with the HAE cultures producing substantially higher levels of RDV-TP that translated into markedly more potent antiviral activity of RDV (**Table 1**). Importantly, the metabolism of RDV in Vero E6 cells appeared altered and was less efficient particularly in comparison with the HAE cultures, indicating that Vero E6 cells might not be an adequate cell type to characterize the antiviral activity of RDV and potentially also other nucleotide prodrug-based antivirals.

**Figure 4:**
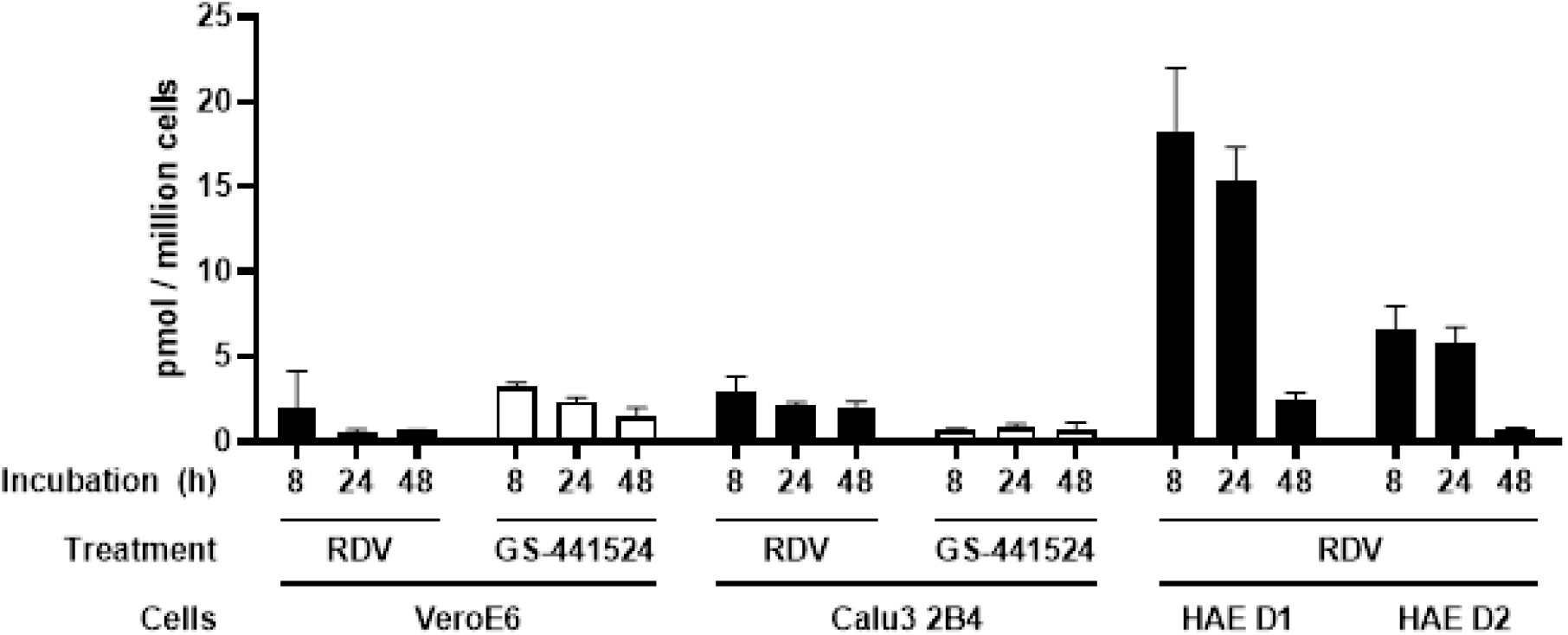
RDV-TP levels in Vero E6, Calu3, and HAE cultures. Vero E6 cells, Calu3 2B4 cells, and HAE cultures were incubated with RDV or GS-441524. At 8, 24, and 48 h of treatment, whole cell extracts were prepared, and RDV-TP levels were quantified by LC-MS/MS as described in Materials and Methods. RDV-TP levels in Vero E6 and Calu3 2B4 cells represent mean ± SD from two independent experiments, each performed with duplicate samples. RDV-TP levels in HAEs represent the mean ± SD of four replicates for each individual donor (D1 and D2).

### RDV is active against the SARS-CoV-2 RdRp *in vivo*

To determine whether RDV exerts antiviral effect on SARS-CoV-2 *in vivo*, we constructed a chimeric mouse-adapted SARS-CoV-1 variant encoding the target of RDV antiviral activity, the RdRp, of SARS-CoV-2 (SARS1/SARS2-RdRp) (**Fig. 5A**). Although other chimeric replicase ORF recombinant CoVs have shown to be viable (Stobart et al., 2013), this is the first demonstration that the RdRp from a related but different CoV can support efficient replication of another. After recovery and sequence-confirmation (**Fig. 5B**) of recombinant chimeric viruses with and without nanoluciferase reporter, we compared SARS-CoV-1 and SARS1/SARS2-RdRp replication and sensitivity to RDV in Huh7 cells. Replication of both viruses was inhibited similarly in a dose-dependent manner by RDV (SARS-CoV-1 mean EC_50_ = 0.007 μM; SARS1/SARS2-RdRp mean EC_50_ = 0.003 μM) (**Fig. 5C and D**). We then sought to determine the therapeutic efficacy of RDV against the SARS1/SARS2-RdRp in mouse models employed for previous studies of RDV (Sheahan et al., 2017). Mice produce a serum esterase absent in humans, carboxyl esterase 1c (Ces1c), that dramatically reduces half-life of RDV. Thus, to mirror pharmacokinetics observed in humans, mouse studies with RDV must be performed in transgenic C57B1/6 *Ces1c^-/-^* mice (Sheahan et al., 2017). We infected female C57B1/6 *Ces1c^-/-^* mice with 10^3^ PFU SARS1/SARS2-RdRp and initiated subcutaneous treatment with 25 mg/kg RDV BID at one day post-infection (dpi). This regimen was continued until study termination. While weight loss did not differ between vehicle- and RDV-treated animals (**Fig. 5E**), lung hemorrhage at five dpi was significantly reduced with RDV treatment (**Fig. 5F**). To gain insight into physiologic metrics of disease severity, we measured pulmonary function daily by whole body plethysmography (WBP). The WBP metric, PenH, is a surrogate marker of pulmonary obstruction (Menachery et al., 2015a). Therapeutic RDV significantly ameliorated loss of pulmonary function observed in the vehicle-treated group (**Fig. 5G**). Importantly, RDV treatment dramatically reduced lung viral load (**Fig. 5H**). Taken together, these data demonstrate that therapeutically administered RDV can reduce virus replication and improve pulmonary function in an ongoing infection with a chimeric SARS-CoV-1/SARS-CoV-2 virus encoding the target of RDV, the RdRp.

**Figure 5.**
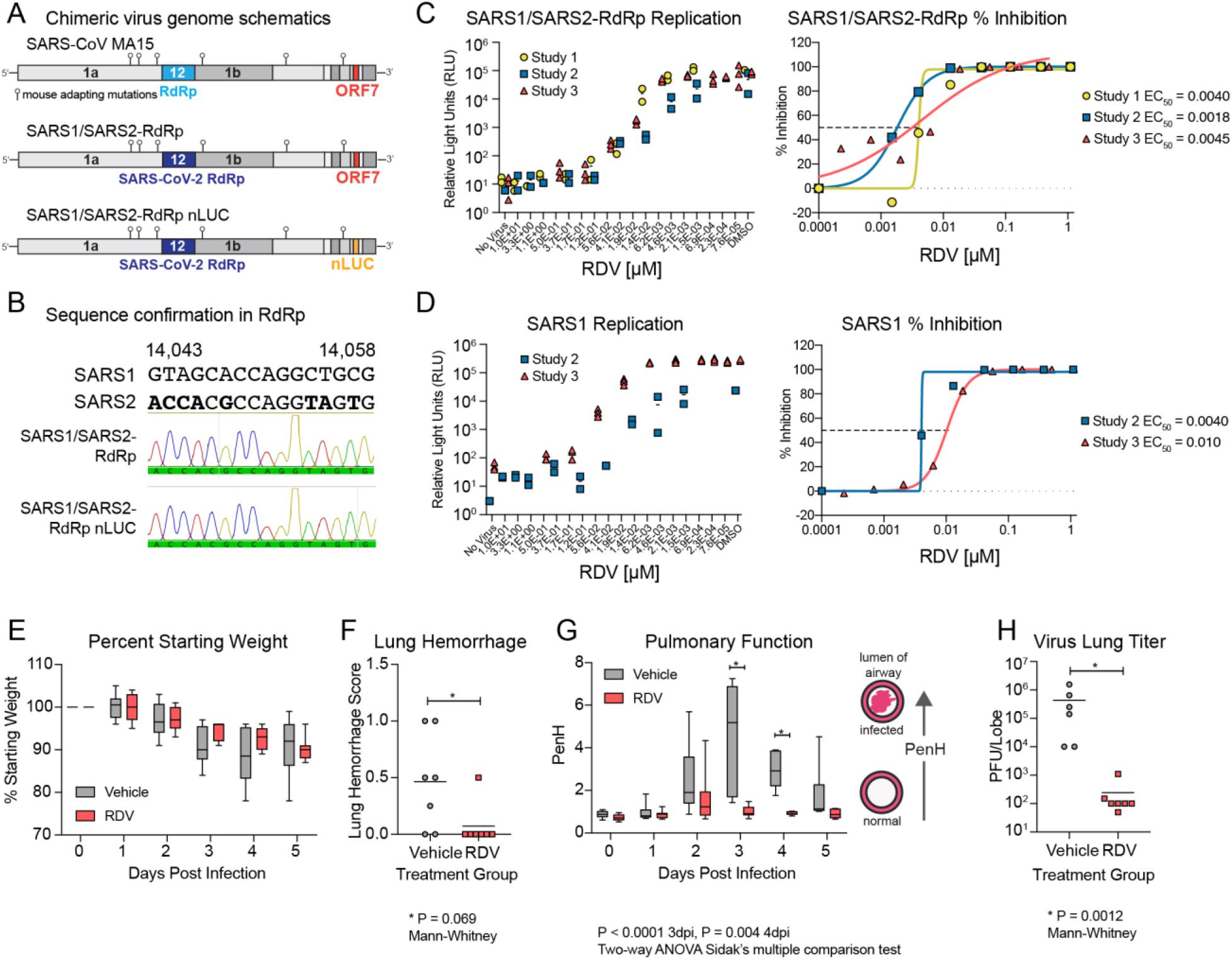
Remdesivir (RDV) is active against the SARS-CoV-2 RdRp *in vivo*. Activity of RDV against the SARS-CoV-2 RdRp was evaluated using a chimeric SARS-CoV-1 encoding the SARS-CoV-2 RdRp (SARS1/SARS2-RdRp). **A.** Schematic of the recombinant SARS-CoV-1 mouse adapted MA15 strain chimeric virus genomes generated for these studies. SARS1/SARS2-RdRp and SARS1/SARS2-RdRp-nLUC were constructed by exchanging the SARS-CoV-1 MA15 RdRp with the SARS-CoV-2 RdRp. ORF7 is replaced by nanoluciferase (nLUC) in SARS2-RdRp-nLUC. **B.** Presence of the SARS-CoV-2 RdRp was confirmed by Sanger sequencing in stocks of both recombinant chimeric viruses. **C.** SARS1/SARS2-RdRp-nLUC replication in Huh7 cells in the presence of RDV (left) and associated percent inhibition (right). **D.** SARS-CoV-1 replication in Huh7 cells in the presence of RDV (left) and associated percent inhibition (right). **E.** Percent starting weight of 17-week old female *Ces1c^-/-^* mice intranasally infected with 1 x 10^3^ PFU of SARS1/SARS2-RdRp and treated with 25mg/kg RDV subcutaneously or vehicle one day post-infection (dpi) and twice daily thereafter. **F.** Lung hemorrhage at five dpi. *P* = 0.069 by Mann-Whitney test. **G.** Pulmonary function by whole-body plethysmography. The PenH metric shown is a surrogate marker of pulmonary obstruction. P < 0.0001 as determined by two-way ANOVA with Sidek’s multiple comparison test. **H.** Lung titer at five dpi as measured by plaque assay. *P* = 0.0012 by Mann-Whitney test. For E, G, boxes encompass 25^th^ to 75^th^ percentile, line is drawn at the median, and whiskers represent the range.

## DISCUSSION

The COVID-19 pandemic has gravely illustrated the need for countermeasures against emerging epidemic and pandemic CoVs. Broad-spectrum antiviral drugs, antibodies, and vaccines are needed to combat the current pandemic and those that will emerge in the future. RDV shows potent activity against an array of genetically diverse CoVs as well as against unrelated emerging viruses like Ebola (Agostini et al., 2018; Brown et al., 2019; Sheahan et al., 2017, 2020a; Warren et al., 2016). In this study, we demonstrate that RDV and its parent nucleoside GS-441524 are active against SARS-CoV-2 in a physiologically relevant cell line and that RDV exerts substantially higher antiviral activity in primary human airway cultures. Potency was directly related to the intracellular concentration of pharmacologically active triphosphate metabolite, which was markedly higher in primary HAE cultures compared to human lung cells (Calu3 2B4) and monkey kidney cells (Vero E6). Our data are consistent with recent studies demonstrating important contributions of natural variation in host and tissue specific gene expression patterns and microbiome specific contributions to drug metabolism, stability, and bioavailability in different tissues (Eriksson, 2013; Koczor et al., 2012). Modeling of RDV onto the SARS-CoV-2 RdRp revealed that the positioning of the RDV into the active site closely resembled that of cognate natural substrate ATP, consistent with efficient incorporation into RNA during replication of the viral genome. RDV decreased viral loads and improved lung function in mice infected with SARS1/SARS2-RdRp chimeric virus when treated at 1 dpi. This is the first rigorous demonstration of potent inhibition of SARS-CoV-2 in continuous and primary human lung cultures and first study suggesting efficacy of RDV against SARS-CoV-2 in mice.

Previous studies of RDV anti-SARS-CoV-2 activity reported EC_50_ values of 0.77 μM as determined by quantification of genome copy number (Wang et al., 2020), 23.15 μM as determined by TCID50, 26.90 as determined by RNA copy number (Choy et al., 2020), and 0.651 μM as determined by cytopathic effect (CPE) (Runfeng et al., 2020), all in Vero E6 cells. The potency of RDV in Vero E6 cells (EC_50_ 1.65 μM) observed in our study is comparable to values reported by Wang *et al*. and Runfeng *et al*., but greater than reported by Choy *et al*. Sequence comparison of the nsp12 from the Seattle, WA isolate used in this study versus SARS-CoV-2 isolates used in the previously mentioned studies assessing RDV potency did not reveal consensus changes in nsp12 sequence, suggesting that any isolate-specific variation in RDV sensitivity is not likely due to differences in the RDV-TP interaction with the RdRp. Therefore, the differences in EC_50_ may be partially explained by intrinsic differences of SARS-CoV-2 virus isolates, quantification methods, and assay conditions such as incubation period and virus input.

Although Vero E6 cells support robust replication of SARS-CoV-2 as illustrated here and elsewhere, our study emphasized the extreme caution that should be exercised when interpreting drug efficacy and potency experiments performed using Vero cell lineages. Nucleoside analog efficacy is greatly dependent on metabolism into the active form. In contrast to GS-441524, RDV contains a protective group which facilitates cellular uptake of the nucleoside analog and addition of the first phosphate group, which accelerates conversion to the active triphosphate (Mehellou et al., 2018). Consistent with previous reports (Agostini et al., 2018), RDV showed enhanced inhibition of SARS-CoV-2 over GS-441524 in Calu3 2B4 cells. In contrast, RDV was two-fold less potent than GS-441524 in Vero E6 cells. Relative potency of the two compounds was directly linked to intracellular concentration of the active triphosphate metabolite, suggesting an altered uptake and/or intracellular metabolism of RDV, consistent with a previous report describing inefficient metabolism of the nucleotide prodrug sofosbuvir in Vero cells (Mumtaz et al., 2017). Drug potency in Vero E6 was similar whether quantified by infectious virus or genome copy number. In Calu3 cells, the potency determined by RT-qPCR was about two-fold lower than when quantified by plaque assay. It is possible that RT-qPCR, which was developed to detect nucleocapsid (N) RNA, also detects packaged subgenomic RNAs and defective genomes in addition to full-length genomes. This would result in underestimation of the reduction in infectious titer. Notably, the potency of RDV against SARS-CoV-1 encoding the SARS-CoV-2 RdRp in Huh7 cells was more than 100-fold higher than that of RDV against bona-fide SARS-CoV-2 in Huh7 and Calu3 cells. This difference could be due to infectivity, which is driven by the SARS-CoV-1 instead of the SARS-CoV-2 spike protein. In addition, SARS-CoV-1 infects Huh7 cells at low frequency at the MOI used in this study and does not appear to spread throughout the culture over the course of the experiment. The number of Huh7 cells in which virus replicates is relatively lower compared to Calu3 cells, possibly enhancing potency of RDV in Huh7 compared to Calu3. Interestingly, the antiviral potency of RDV against SARS-CoV-2 in HAE cultures was comparable to SARS-CoV-1 and MERS-CoV (Sheahan et al., 2017), which is consistent with the high conservation of the RdRp active site across these different CoVs. Together, these results emphasize the need for careful selection and use of multiple cell types and methods to study potency and efficacy of nucleoside analogs and other antiviral compounds.

The target of RDV antiviral activity is the viral RdRp. To mirror the pharmacokinetic exposures observed in humans, RDV studies in mice must be performed in *Ces1c-/-* animals (Sheahan et al., 2017). In addition, SARS-CoV-2 does not readily infect WT mice due to incompatibilities between virus spike and the murine ortholog of ACE2, which serves as the SARS-CoV-2 entry receptor (Wan et al., 2020). The breeding a doubly transgenic (hACE2, *Ces1c-/-*) mice for use in RDV efficacy studies is ongoing. To rapidly assess the therapeutic efficacy of RDV against SARS-CoV-2, we constructed recombinant SARS-CoV-1 chimeric virus encoding the SARS-CoV-2 RdRp (SARS1/SARS2-RdRp). Virus entry, tropism, and pathogenesis of this chimeric virus are driven by parental mouse-adapted SARS-CoV-1 virus. Similar to our previous studies with SARS-CoV-1 (Sheahan et al., 2017), we now show that therapeutic administration of RDV one dpi can both reduce viral load and improve pulmonary function in mice. The kinetics of SARS-CoV-1 replication and disease are notably compressed in mice as compared to humans where virus titer peaks 10-15 days after onset of symptoms (Hung et al., 2004; Peiris et al., 2004). By comparison, initial reports suggest that SARS-CoV-2 replication peaks around 5-6 days after symptom onset, just prior to onset of dyspnea (Pan et al., 2020; Zhou et al., 2020a). A recent preprint described the therapeutic efficacy of RDV against SARS-CoV-2 in rhesus macaques, where RDV treatment reduced respiratory pathology and viral loads in bronchoalveolar lavage fluid (Williamson et al., 2020). Prior to the emergence of pandemic SARS-CoV-2, RDV was evaluated in phase 1 clinical trials as well as phase 2 randomized controlled trial trials to treat acute Ebola virus disease in the Democratic Republic of Congo (DRC), and human safety data are available (Mulangu et al., 2019). Thus, our preclinical development of RDV positioned RDV for immediate compassionate use of RDV for severely ill COVID-19 patients. While early results are promising (Grein et al., 2020), Phase III randomized controlled trials for the treatment of patients with COVID-19 are ongoing globally and will ultimately determine efficacy, safety, and optimal dosing of RDV in patients with different stages of COVID-19.

Despite worldwide drug discovery efforts and over 300 active clinical evaluations of potential treatments, no effective countermeasure currently exists to combat COVID-19 (Sanders et al., 2020) or likely future CoV pandemics. Large-scale deployment of antiviral monotherapies creates high risk for emergence of drug resistance. Our previous work demonstrates that CoV resistance to RDV is generated slowly and is conferred by two mutations in the RdRp. In addition, RDV-resistant CoVs exhibit reduced replication capacity and are also more sensitive to another potently active nucleoside analog inhibitor β-D-*N*4-hydroxycytidine (NHC; EIDD-1931/2801) (Agostini et al., 2019; Sheahan et al., 2020b). Therapies combining direct-acting antivirals (DAAs) such as RDV and NHC, along with other DAAs such as antibodies and protease inhibitors that target different stages of the viral replication cycle, could be considered for counteracting resistance if it emerges in patients treated with antiviral monotherapy. In addition, attention should be given to combining DAAs with anti-inflammatory drugs to potentially extend the treatment window during which DAAs can improve outcomes. With SARS-CoV-1-, SARS-CoV-2-, and MERS-like CoVs continuing to circulate in bat species, more outbreaks of novel CoVs are expected (Menachery et al., 2015b, 2016). Identification and evaluation of broadly efficacious, robust anti-CoV therapies are thus urgently needed in the present and future.

## ACKNOWLEDGEMENTS

This project was funded in part by the National Institute of Allergy and Infectious Diseases, National Institutes of Health, Department of Health and Human Service awards: 1U19AI142759 (Antiviral Drug Discovery and Development Center awarded to M.R.D. and R.S.B); 5R01AI132178 awarded to T.P.S. and R.S.B.; and 5R01AI108197 awarded to M.R.D. and R.S.B. D.R.M was funded by T32 AI007151 and a Burroughs Wellcome Fund Postdoctoral Enrichment Program Award. The Marsico Lung Institute Tissue Procurement and Cell Culture Core is supported by NIH grant DK065988 and Cystic Fibrosis Foundation grant BOUCHE15RO. We also are grateful for support from the Dolly Parton COVID-19 Research Fund, the VUMC Office of Research, and the Elizabeth B. Lamb Center for Pediatric Research at Vanderbilt University.

We thank Dr. Natalie Thornburg at the Centers for Disease Control and Prevention in Atlanta, USA for providing the stock of SARS-CoV-2 used in this study. Finally, we thank VUMC and UNC Environmental Health and Safety personnel for ensuring that our work is performed safely and securely. We also thank Facilities Management personnel for their tireless commitment to excellent facility performance and our grant management teams for their administrative support of our research operations.

## AUTHOR CONTRIBUTIONS

A.J.P., T.P.S., conceived, designed, and performed experiments and management and coordinated responsibility for the research activity planning and execution. A.J.P., J.K.P., J.P.B, and T.P.S. wrote the manuscript. A.S.G., A.S., S.R.L, K.H.D., B.L.Y., M.L.A., L.J.S., J.D.C., X.L., T.M.H., K.G., D.R.M., A.J.B., R.L.G., J.K.P., V.D.P., J.P., B.M., D.B., and E.M. performed experiments. T.P.S, D.R.M, R.S.B., A.J.P., J.D.C., and M.R.D. secured funding. D.P.P, N.J.T, T.C. provided reagents. J.D.C., J.Y.F., J.P.B., D.P.P, T.C., R.S.B., M.R.D edited the manuscript and provided expertise and feedback.

## METHODS

### Cells, viruses, and compounds

Vero (ATCC CCL-81) and Vero E6 (ATCC CRL-1586) cells were purchased from ATCC and cultured in DMEM supplemented with 10% fetal bovine serum (FBS) (Gibco, ThermoFisher Scientific) or 10% FCS fetal clonal serum (FCS)(HyClone, GE Life Sciences), 100 U/ml penicillin and streptomycin (Gibco, ThermoFisher Scientific), and 0.25 μM amphotericin B (Corning). Human hepatoma (Huh7) cells were provided by Dr. Mark Heise at UNC Chapel Hill and grown in DMEM supplemented with 10% FBS (Hyclone) and 1× antibiotic-antimycotic (Gibco, ThermoFisher Scientific). Calu3 2B4 cells (Yoshikawa et al., 2010) were cultured in DMEM supplemented with 20% FBS, 100 U/mL penicillin and streptomycin, and 0.25 μM amphotericin B.

Primary HAE cell cultures used in antiviral activity assays were obtained from the Tissue Procurement and Cell Culture Core Laboratory in the Marsico Lung Institute/Cystic Fibrosis Research Center at UNC. All assays in this report were performed using a single HAE cell donor. Human tracheobronchial epithelial cells provided by Dr. Scott Randell were obtained from airway specimens resected from patients undergoing surgery under University of North Carolina Institutional Review Board-approved protocols (#03-1396) by the Cystic Fibrosis Center Tissue Culture Core. Primary cells were expanded to generate passage 1 cells and passage 2 cells were plated at a density of 250,000 cells per well on Transwell-COL (12mm diameter) supports (Corning). Human airway epithelium cultures (HAE) were generated by provision of an air-liquid interface for 6 to 8 weeks to form well-differentiated, polarized cultures that resembled in vivo pseudostratified mucociliary epithelium (Fulcher et al., 2005).

Clinical specimens of SARS-CoV-2 from a case-patient who acquired COVID-19 during travel to China and diagnosed in Washington State, USA upon return were collected as described (Holshue et al., 2020). Virus isolation from this patient’s specimens was performed as described in (Harcourt et al.). The sequence is available through GenBank (accession number MN985325). A passage 3 stock of the SARS-CoV-2 Seattle isolate was obtained from the CDC and passed twice in Vero E6 cells to generate high-titer passage 5 stock for experiments described in this manuscript.

SARS-CoV-1 expressing GFP (GFP replaces ORF7) was created from molecular cDNA clones as described (Scobey et al., 2013; Sims et al., 2005). To create SARS-CoV-1 expressing nanoluciferase (nLUC), the gene for GFP was replaced with nLUC and isolated using our existing mouse adapted SARS-CoV-1 (MA15) SARS-CoV-1 Urbani molecular clone (Yount et al., 2003). A synthetic cDNA encoding the SARS-CoV-2 RdRp (Integrated DNA Technologies) was cloned into SARS MA15 D fragment using StuI (5’) and BsaI (3’) via Gibson assembly. The resultant plasmids were sequence confirmed and then utilized to generate recombinant virus with or without nanoluciferase as described. Recombinant virus stocks were confirmed to harbor SARS-CoV-2 RdRp by Sanger sequencing.

Remdesivir (RDV; GS-5734) and GS-441524 were synthesized by the Department of Medicinal Chemistry, Gilead Sciences (Foster City, CA).

### Modeling

A model of the elongating SARS-CoV-2 polymerase complex was generated based on a homology model which used the cryo-EM structure of apo SARS-CoV-1 as a template (PDB: 6NUR, (Kirchdoerfer and Ward, 2019) as described previously (Gordon et al., 2020b). RNA primer and template, a substrate ATP and two catalytic Mg^++^ ions were oriented in the structure based on alignment to a ternary x-ray structure of HCV NS5B (PDB: 4WTD, (Appleby et al., 2015). The structure was then optimized with a series of constrained minimizations. To this structure we aligned the new cryo-EM structure of the SARS-CoV-2 replication complex. As the SARS-CoV-2 structure does not significantly differ from the SARS-CoV-1 structure, rather than a complete replacement of the model, we incorporated only those residues that had not been resolved in the previous structure (residues 31-116). Additional optimization, particularly of the RNA in the exit channel, was done following the previously outlined procedures. After optimization of the ATP structure, RDV-TP was modeled into the active site and minimized. Models of the V557L and F480L mutants and the other coronavirus models reported here were generated based on this final model. All work was carried out using Prime and Macromodel (Schrödinger, LLC, New York, NY, 2020). 3D coordinates of the SARS-CoV-2/RDV-TP model are provided in the Supplementary Material.

### Sequence alignments

Coronavirus nsp12 sequence alignment was generated using CLC Workbench (Qiagen) from sequences downloaded from the NCBI website (Accession numbers MN985325.1 and MT123290.1) and from GISAID’s EpiFlu™ Database (Elbe and Buckland-Merrett, 2017; Shu and McCauley, 2017): Accession ID EPI_ISL_402124; virus name hCoV-19/Wuhan/WIV04/2019; Location: Asia / China / Hubei / Wuhan; Collection date 2019-12-30 Originating lab Wuhan Jinyintan Hospital; Submitting lab Wuhan Institute of Virology, Chinese Academy of Sciences; Authors Peng Zhou, Xing-Lou Yang, Ding-Yu Zhang, Lei Zhang, Yan Zhu, Hao-Rui Si, Zhengli Shi. Accession ID EPI_ISL_412028; virus name hCoV-19/Hong Kong/VM20001061/2020; Location Asia / Hong Kong; Collection date 2020-01-22; Originating lab Hong Kong Department of Health; Submitting lab: School of Public Health, The University of Hong Kong; Authors Dominic N.C. Tsang, Daniel K.W. Chu, Leo L.M. Poon, Malik Peiris.

### Replication in different cell types

Vero E6, Vero, Calu3 2B4, and Huh7 cells were seeded in 24 well plates and allowed to adhere for 24 h. Cells were adsorbed with 100 μl SARS-CoV-2 in gel saline for 30 minutes (min) at 37°C with manual rocking every 10 min. Virus inoculum was removed, cells were washed in PBS, and 0.5 ml medium was added to each well. Supernatant was collected at 0, 24, 48, and 72 h post-infection, and infectious viral titer in supernatants was determined by plaque assay.

### Antiviral activity assays

Vero E6 cells were seeded at 1 x 10^5^ cells per well, and Calu3 2B4 cells were seeded at 2 x 10^5^ cells per well in 24-well plates (Corning). Cells were allowed to adhere for 16-24 h. Drugs were dissolved in DSMO and serially diluted in DMSO to achieve 1000x final concentration. Equal volumes of each 1000x concentration were further diluted 1000-fold in medium up to 2 h before start of the infection. Cells were adsorbed at MOI = 0.01 PFU/cell with SARS-CoV-2 in gel saline for 30 min at 37°C. Plates were rocked manually to redistribute the inoculum every 10 minutes. Viral inoculum was removed, and cells were washed with pre-warmed PBS+/+ (Corning) for 5 minutes. PBS+/+ was removed, and medium containing dilutions of RDV, GS-441524, or vehicle (DMSO) was added. Cells were incubated at 37°C. At 48 (Vero E6) or 72 (Calu3 2B4) hpi, supernatants were harvested and processed for plaque assay and RT-qPCR.

Huh7 cells were plated at a density of 8 x 10^4^ cells per well. Twenty-four hours later, fresh medium was added. Triplicate wells of cells were infected for 1 h at 37°C with SARS1/SARS2-RdRp-nLUC or SARS-CoV-1-nLUC diluted 1:100 in culture medium. Virus was removed, cultures were rinsed once with medium, and fresh medium containing dilutions of RDV or vehicle (DMSO) was added. DMSO (0.05%) was constant in all conditions. At 48 hpi, virus replication was measured by nLUC assay (Promega) using a SpectraMax plate reader (Molecular Devices).

Before infection, HAE cultures (approximately 1 x 10^6^ cells per well) were washed with phosphate-buffered saline (PBS) and moved into air-liquid interface medium containing various doses of RDV ranging from 0.00260 to10 mM (final DMSO, <0.05%). Cultures were infected with SARS-CoV-2 clinical isolate (2019-nCoV/USA-WA1/2020) at MOI = 0.5 PFU/cell for 2 h at 37°C, after which virus was removed and cultures were washed three times with PBS, followed by incubation at 37°C for 48 h. The apical surface of each culture was washed with PBS and collected for virus titration, measured as plaque-forming units (PFU) as previously described for SARS-CoV-1 (Scobey et al., 2013; Sims et al., 2005).

### Cytotoxicity Assays

Cells were seeded at a density of 15,000 cells/well (Vero E6) or 30,000 cells/well (Calu3 2B4) in a white-wall clear-bottom 96-well plate (Corning) and incubated at 37°C overnight. Medium was removed, and serial dilutions of drug in medium were added to each well (see “**antiviral activity assays**”). Cytotoxicity was determined using CellTiterGlo Cell Viability Assay (Promega) according to manufacturer’s instructions at 48 h (Vero E6) or 72 h (Calu3). HAE cultures were treated with the same concentration range of drug in Transwell plates (Corning). Cytotoxicity in HAE was previously determined by RT-qPCR of TRIzol-extracted RNA (Sheahan et al., 2017)

### Quantification of infectious viral titer by plaque assay

Approximately 1 x 10^6^ Vero E6 cells/well were seeded in 6-well tissue culture plates (Corning) and allowed to grow to confluence for 48 h. Medium was removed, and 200 μL of 10-fold serial dilutions of virus-containing supernatants in gel saline were adsorbed in duplicate for 30 min at 37°C. Plates were rocked manually to redistribute inoculum every 10 minutes. Cells were overlaid with a 1:1 mixture of 2x DMEM and 2% agar in ddH2O and incubated at 37°C. Plaques were enumerated in unstained monolayers at 48-72 hpi using a light box.

### Quantification of viral RNA genome copy number by RT-qPCR

Cell supernatants were harvested in TRIzol LS reagent (Invitrogen), and RNA was purified following phase separation by chloroform as recommended by the manufacturer. RNA in the aqueous phase was collected and further purified using PureLink RNA Mini Kits (Invitrogen) according to manufacturer’s protocol. Viral RNA was quantified by reverse-transcription quantitative PCR (RT-qPCR) on a StepOnePlus Real-Time PCR System (Applied Biosystems) using TaqMan Fast Virus 1-Step Master Mix chemistry (Applied Biosystems). SARS-CoV-2 N gene RNA was amplified using forward (5’-GACCCCAAAATCAGCGAAAT) and reverse (5’-TCTGGTTACTGCCAGTTGAATCTG) primers and probe (5’-FAM-ACCCCGCATTACGTTTGGTGGACC-BHQ1) designed by the United States Centers for Disease Control and Prevention (oligonucleotides produced by IDT, cat# 10006606). RNA copy numbers were interpolated from a standard curve produced with serial 10-fold dilutions of N gene RNA. Briefly, SARS-CoV-2 N gene positive control plasmid (IDT, cat# 10006625) served as template to PCR-amplify a 1280 bp product using forward (5’-TAATACGACTCACTATAGGGATGTCTGATAATGGACCCCA) and reverse (5’-TTAGGCCTGAGTTGAGTCAG) primers that appended a T7 RNA polymerase promoter to the 5’ end of the complete N ORF. PCR product was column purified (Promega) for subsequent *in vitro* transcription of N RNA using mMESSAGE mMACHINE T7 Transcription Kit (Invitrogen) according to manufacturer’s protocol. N RNA was purified using RNeasy mini kit (Qiagen) according to manufacturer’s protocol, and copy number was calculated using SciencePrimer.com cop number calculator.

### *In vitro* metabolism of RDV and GS-441524

Calu3 2B4 or Vero E6 cells were seeded in a 6-well plate at 8.0 x 10^5^ or 3.5 × 10^5^ cells/well, respectively. Twenty-four hours later, cell culture media was replaced with media containing 1 μM RDV (GS-5734) or GS-441524 and incubated at 37°C. Differentiated HAE cultures from two healthy donors (MatTek Corporation; Ashland, MA) were maintained with media replacement every other day for 1 week. The HAE donors were 56- and 62-year-old females of the same race. At the time of treatment, media was replaced on the basal side of the transwell HAE culture, while the apical surface media was replaced with 200 μL media containing 1 μM RDV. At 8, 24 and 48h post drug addition to all cultures, cells were washed 3 times with ice-cold tris-buffered saline, scraped into 0.5 mL ice-cold 70% methanol and stored at −80°C. Extracts were centrifuged at 15,000 x g for 15 minutes and supernatants were transferred to clean tubes for evaporation in a MiVac Duo concentrator (Genevac). Dried samples were reconstituted in mobile phase A containing 3 mM ammonium formate (pH 5) with 10 mM dimethylhexylamine (DMH) in water for analysis by LC-MS/MS, using a multi-stage linear gradient from 10% to 50% acetonitrile in mobile phase A at a flow rate of 300 μL/min. Analytes were separated using a 50 x 2 mm, 2.5 μm Luna C18(2) HST column (Phenomenex) connected to an LC-20ADXR (Shimadzu) ternary pump system and HTS PAL autosampler (LEAP Technologies). Detection was performed on a Qtrap 6500+ (AB Sciex) mass spectrometer operating in positive ion and multiple reaction monitoring modes. Analytes were quantified using a 7-point standard curve ranging in concentration from 0.156 to 40 pmol prepared in extracts from untreated cells. For normalization by cell number, multiple untreated Calu3 or Vero E6 culture wells were counted at each timepoint. HAE cells were counted at the 24-h timepoint and the counts for other timepoints were determined by normalized to endogenous ATP levels for accuracy.

### Formulations for *in vivo* studies

RDV was solubilized at 2.5 mg/mL in vehicle containing 12% sulfobutylether-β-cyclodextrin sodium salt in water (with HCl/NaOH) at pH 5.0.

### *In vivo* efficacy studies

All animal experiments were performed in accordance with the University of North Carolina at Chapel Hill Institutional Animal Care and Use Committee policies and guidelines. To achieve a pharmacokinetic profile similar to that observed in humans, we performed therapeutic efficacy studies in *Ces1^-/-^* mice (stock 014096, The Jackson Laboratory), which lack a serum esterase not present in humans that dramatically reduces RDV half-life (Sheahan et al., 2017). 17 week-old female *Ces1^-/-^* mice were anaesthetized with a mixture of ketamine/xylazine and intranasally infected with 10^3^ PFU SARS1/SARS2-RdRp in 50 μL. One dpi, vehicle (n = 7) and RDV (n = 7) dosing was initiated (25 mg/kg subcutaneously) and continued every 12 h until the end of the study at five dpi. To monitor morbidity, mice were weighed daily. Pulmonary function testing was performed daily by whole body plethysmography (WBP) (Data Sciences International) (Sheahan et al., 2017). At five dpi, animals were sacrificed by isoflurane overdose, lungs were scored for lung hemorrhage, and the inferior right lobe was frozen at −80°C for viral titration via plaque assay on Vero E6 cells. Lung hemorrhage is a gross pathological phenotype readily observed by the naked eye and driven by the degree of virus replication, where lung coloration changes from pink to dark red (Sheahan et al., 2017, 2020a). For the plaque assay, 5 x 10^5^ Vero E6 cells/well were seeded in 6-well plates. The following day, medium was removed, and monolayers were adsorbed at 37°C for one h with serial dilutions of sample ranging from 10^-1^ to 10^-6^. Cells were overlayed with 1X DMEM, 5% Fetal Clone 2 serum, 1× antibiotic-antimycotic, 0.8% agarose. Viral plaques were enumerated three days later.

### Mathematical and statistical analyses

The EC_50_ value was defined in GraphPad Prism 8 as the concentration at which there was a 50% decrease in viral replication relative to vehicle alone (0% inhibition). Curves were fitted based on four parameter non-linear regression analysis. All statistical tests were executed using GraphPad Prism 8.

