## Supplemental materials for "Remdesivir potently inhibits SARS-CoV-2 in human lung cells and chimeric SARS-CoV expressing the SARS-CoV-2 RNA polymerase in mice"

### Supplemental Figure 1.

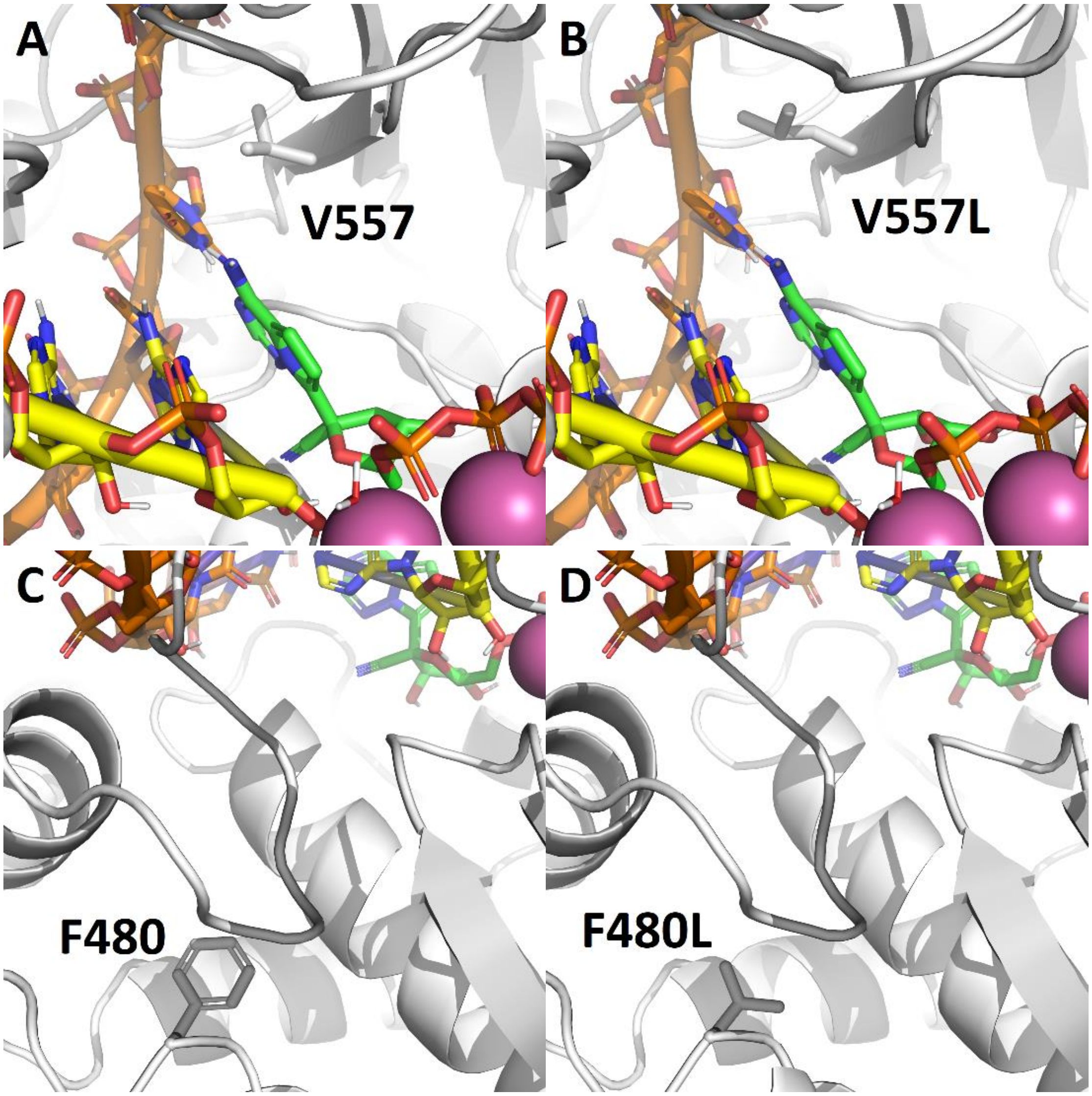


**Figure S1. A.** Influence of RDV resistance mutations in MHV selected by virus passage in the presence drug. WT V557 is in direct contact with the template base. **B.** V557L leads to a modest repositioning of the template, and by extension, RDV (green). **C.** WT F480 lies outside of the active site. **D.** F480L leads to minor adjustments in structural elements that form both the active site and RNA binding pocket.

### Supplemental Figure 2.

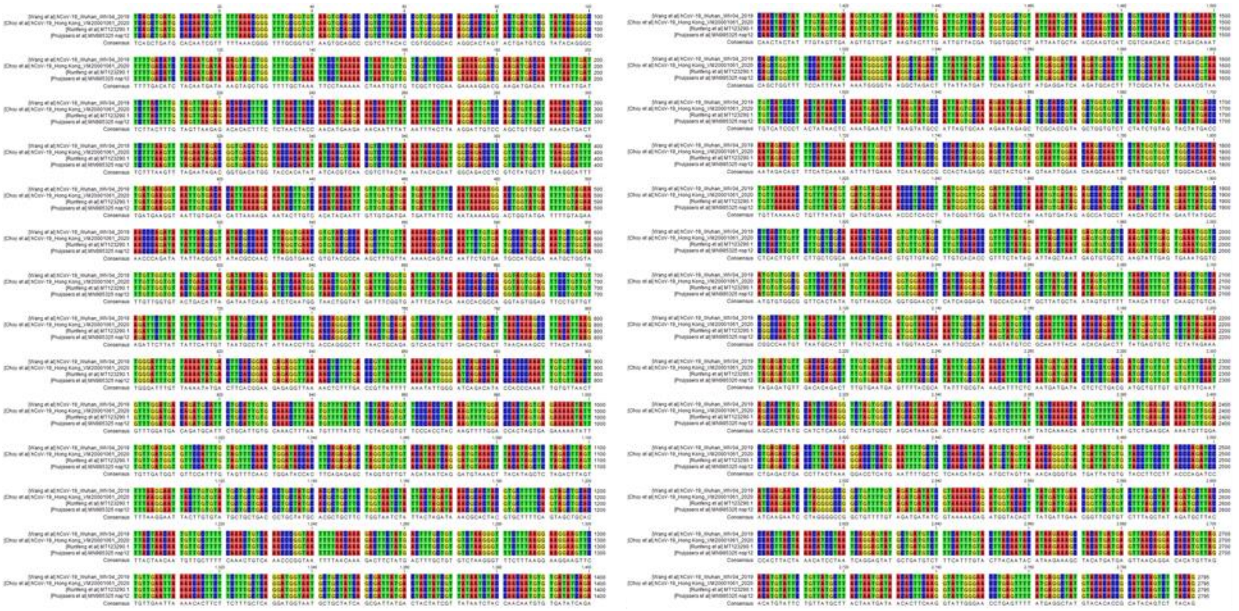

**Figure S2. Nucleotide sequence conservation of *nsp12* from different SARS-CoV-2 isolates in RDV studies.** Alignment of full nsp12 nucleotide sequences from isolates hCoV-19_Wuhan_WIV04_2019 (GISAID EpiFlu™ Database Accession ID: EPI_ISL_402124), hCoV-19_Hong Kong_VM20001061_2020 (GISAID EpiFlu™ Database Accession ID: EPI_ISL_412028), SARS-CoV-2/human/CHN/IQTC01/2020 (GenBank Accession number MT123290.1), and 2019-nCoV/USA-WA1/2020 (GenBank Accession number MN985325.1). The nsp12 sequences are 100% conserved at the nucleotide level.

### Supplemental Figure 3.

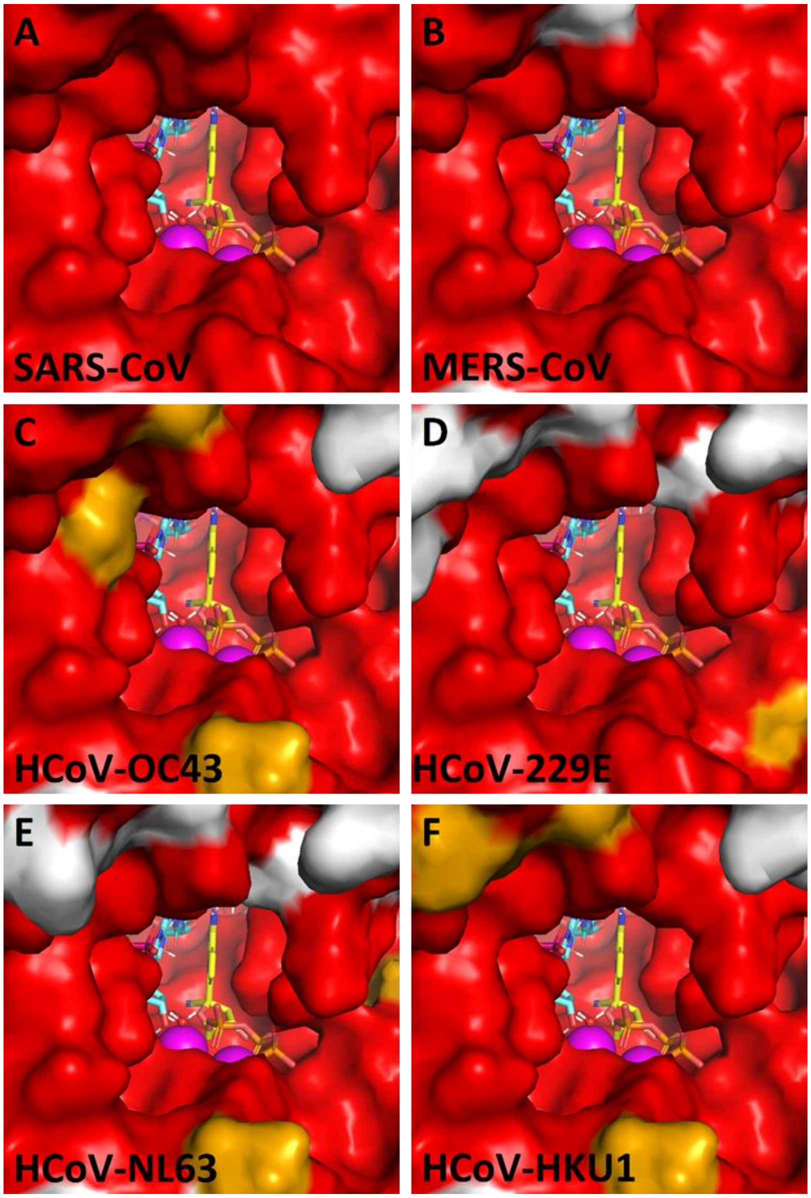


**Figure S3. Models of RDV-TP in other human coronaviruses. A.** SARS-CoV [AAP13442.1] **B.** MERS-CoV [AFS88944.1] **C.** HCoV-OC43 [AAX85675.1] **D.** HCoV-229E [AFR79248.1] **E.** HCoV-NL63 [AFV53147.1] **F.** HCoV-HKU1 [ABD75567.1]. Residues in red are conserved relative to SARS-CoV-2 [QHD43415.1]. Residues in gold are similar. Residues in white are dissimilar. SARS-CoV-2 is identical to SARS-CoV-1 out to a radius of 18 Å from the active site. While differences are visible on the periphery of the active site, residues that interact directly with the RDV-TP are highly conserved for all human CoVs.

### Supplemental Table 1: Metabolite levels following RDV or GS-441524 treatment of Vero E6 and Calu3 cell lines

| Treatment | Time (h) | Metabolite levels (pmol / million cells)[a, b] | | | | | | | |
|---|---|---|---|---|---|---|---|---|---|
| | | RDV-TP | | RDV-DP | | RDV-MP | | GS-441524 | |
| | | VeroE6 | Calu3 | VeroE6 | Calu3 | VeroE6 | Calu3 | VeroE6 | Calu3 |
| RDV | 8 | 2.02 ± 1.87[c] | 2.87 ± 0.84 | BLQ | 2.90 ± 2.03 | BLQ | 2.43 ± 2.14 | BLQ | 0.95 ± 0.20 |
| | 24 | 0.54 ± 0.15 | 2.17 ± 0.14 | BLQ | 1.12 ± 0.11 | BLQ | 0.50 ± 0.03 | BLQ | 0.64 ± 0.04 |
| | 48 | 0.67 ± 0.03 | 2.00 ± 0.31 | BLQ | 1.13 ± 0.03 | BLQ | 0.31 ± 0.06 | BLQ | 1.02 ± 0.07 |
| GS-441524 | 8 | 3.19 ± 0.25 | 0.67 ± 0.09 | BLQ | 0.35 ± 0.04 | BLQ | 0.06 ± 0.02 | 2.67 ± 0.72 | 2.96 ± 0.59 |
| | 24 | 2.37 ± 0.22 | 0.85 ± 0.16 | 0.35 ± 0.05 | 0.73 ± 0.45 | BLQ | 0.08 ± 0.04 | 1.45 ± 0.46 | 3.56 ± 0.46 |
| | 48 | 1.49 ± 0.44 | 0.72 ± 0.34 | 0.23 ± 0.03 | 0.72 ± 0.16 | BLQ | 0.13 ± 0.07 | 1.31 ± 0.70 | 4.31 ± 0.60 |

TP: triphosphate; DP: diphosphate; MP: monophosphate.

[a] Values represent mean ± SD from two independent experiments, each performed with duplicate samples.

[b] BLQ (below limit of quantitation). Limit of quantification was as follows for each analyte: RDV-DP, 0.156 pmol/sample; RDV-MP, 0.039 pmol/sample; GS-441524, 0.625 pmol/sample

[c] Analysis of one technical replicate at 8 h following treatment of Vero E6 cells had a markedly high RDV-TP level compared to three other replicates. All four values are included in the mean +/- SD calculations.

### Supplemental Table 2: Metabolite levels following RDV treatment of primary HAE cultures

| Donor[a] | Time (h) | Remdesivir metabolite levels (pmol / million cells)[b,c] | | | |
|---|---|---|---|---|---|
| | | RDV-TP | RDV-DP | RDV-MP | GS-441524 |
| 1 | 8 | 18.3 ± 3.22 | 2.10 ± 0.14 | 0.54 ± 0.10 | BLQ |
| | 24 | 15.3 ± 1.73 | 3.45 ± 0.46 | 1.31 ± 0.32 | BLQ |
| | 48 | 2.45 ± 0.36 | BLQ | BLQ | BLQ |
| 2 | 8 | 6.58 ± 1.18 | 0.87 ± 0.13 | 0.57 ± 0.15 | BLQ |
| | 24 | 5.78 ± 0.84 | 1.72 ± 0.28 | 1.19 ± 0.20 | BLQ |
| | 48 | 0.73 ± 0.07 | BLQ | BLQ | BLQ |

[a] Origin of tissues are from healthy, non-smoker donors. Donor 1 = 56-year-old black female; Donor 2 = 62-year-old black female

[b] Values represent mean ± SD from four independent replicates for each donor

[c] BLQ (below limit of quantitation); Limit of quantification for each analyte is as follows: RDV-DP, 0.156 pmol/sample; RDV-MP, 0.156 pmol/sample; GS-441524, 0.625 pmol/sample

### Supplemental Figure 4.

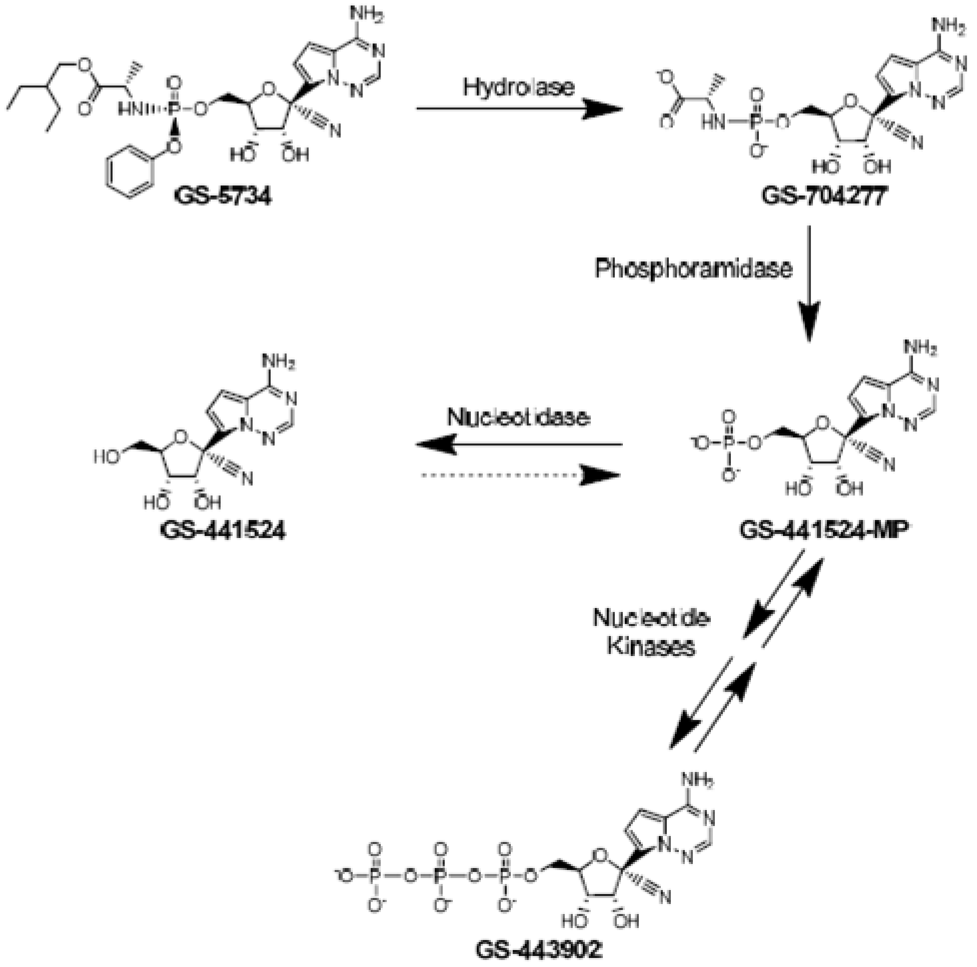


**Figure S4: Generalized intracellular metabolic pathway of remdesivir.** Combined results from pharmacology and pharmacokinetic studies have led to the proposed intracellular metabolic pathway. Remdesivir (GS.5734) is activated to the pharmacologically active nucleoside analog triphosphate, GS-443902, by a sequential metabolic activation pathway. A cellular hydrolase removes the ester, then a spontaneous chemical step forms the intermediate metabolite GS-704277. Phosphoramidase activity subsequently cleaves the phosphoramide bond, liberating the nucleoside analog monophosphate (GS-441524-MP). GS-441524-MP is either catalyzed to the active triphosphate, GS-443902, by nucleotide kinases or dephosphorylated to the nucleoside analog GS-441524.
